# A fast and easy mass sampling method for social bee crop content

**DOI:** 10.64898/2026.01.27.701953

**Authors:** Laura Degirmenci, Martin Gabel, Jessica Strehl, Ricarda Scheiner

## Abstract

Honey bees (*Apis mellifera*) are capable of foraging over large distances around their nest and can thus serve as ideal bioindicators for various research questions. While foraging for nectar, they collect a variety of pollutants such as environmental toxins and agrochemicals, which can be traced in their digestive tract. Likewise, naturally occurring compounds in a bee’s crop content (e.g., different sugars and pollen grains) can be analyzed to assess the foraging preferences of bees or to study the species composition of flowering plants in the respective environment. Regardless of the research question, a precise, reproducible, and efficient method for examining pollinators’ crop contents is needed to fully exploit the honey bee’s potential as a bioindicator. Existing methods are often insufficiently precise with respect to the recovered volume and content, too time-consuming or lack the opportunity to store samples flexibly. Consequently, they do not meet the requirements of large-scale sampling commonly needed in scientific field work. We present a novel, rapid and scalable method for fast and easy mass sampling of honey bee crop contents which holds potential for adaptation to other pollinator taxa such as bumblebees. With this method individuals can be collected in the field and stored prior to sample processing. Crop contents can be obtained later on by centrifuging the entire individual, thereby decoupling sample collection from further analysis. Afterwards, insect samples and extracted fluids can be stored again, enabling subsequent analysis in an efficient and time-saving manner. We demonstrate that crop contents of previously fed animals can be obtained precisely in terms of both volume and composition and present an example of application in honey bee foraging ecology as a proof of concept.

## Introduction

Honey bees (*Apis mellifera*) are generalist foragers that exploit a wide variety of flowering plants as food resources (Winston, 1991; Seeley, 2014). They are important key pollinators in both natural and agricultural ecosystems (Meixner, 2010; Potts, et al., 2010; Garibaldi, et al., 2011). When foraging for nectar and pollen, honey bees are able to evaluate, learn and communicate the location and quality of their food source (Scheiner, et al., 2020; George, et al., 2020; Finke, et al., 2021). As social insects comprising thousands of individuals, honey bees transport large quantities of nectar and pollen into their nest and can forage within a radius exceeding 3 km (Beekman and Ratnieks, 2000), covering a wide range of plant species (Winston, 1991; Seeley, 2014). Therefore, the crop contents of honey bee foragers, i.e., the liquids collected outside the nest, are of particular interest for various research fields. This includes studies on the foraging behavior of this readily available model organism, as well as related studies on pollination ecology (Hung, et al., 2018) and division of labor (Page, et al., 2006).

Besides their importance as a model organisms, honey bees can serve as bioindicators: their crop contents can be analyzed to quantify environmental pollution (Celli & Maccagnani, 2003). For example, honey bees and hive products can be screened for environmental toxins such as agrochemicals, or traffic-related pollutants to assess exposure levels in the surrounding environment (Herrero-Latorre et al., 2017; Costa, et al., 2019; Goretti, et al., 2020; Di Fiore, et al., 2022). Honey bee crop contents thus provide valuable samples for long-term monitoring approaches, e.g., when focusing on environmental pollution based on contaminants or changes in floral resources and plant species composition based on pollen and sugar spectra.

For more detailed investigations, it may also be useful to include crop contents of other pollinators occupying different ecological niches or exhibiting greater specialization in foraging or nesting ecology. For instance, some bumblebee species (e.g., the buff-tailed bumblebee *Bombus terrestris)* are commonly bred for agricultural pollination and therefore easily accessible and manageable under controlled conditions (Röseler, 1985; Goulson, 2003). Similarly, the mason bee species *Osmia bicornis* and *Osmia cornuta*, are frequently used as study organisms and thus represent promising candidates for an extension of crop content analysis in future research (Brandt, et al., 2020). Although honey bees, mason bees and bumblebees partly overlap in their food resources, they nevertheless show distinct differences in their nesting biology and activity periods (Lyu, et al., 2023). Thus, detailed investigations of their crop contents may help to gain deeper insights into all of the above-mentioned study fields.

Numerous methods have been developed to collect liquids from the honey bee’s crop. In general, these methods either rely on dissecting the honey sacs or manually squeezing crop contents through the esophagus, as summarized by Gary & Lorenzen (1976) and Sylvester et al. (1983). Both approaches require labor-intensive and time-consuming handling of individual foragers.

Additionally, portions of the crop contents are often lost, because dying bees regurgitate, leading to unusable samples or unreliable measurements. More recently, Reetz and Wallner (2014) described an advanced technique involving the preparation of frozen honey bees’ abdominal segments in order to centrifuge the crop contents. The collection and investigation of crop contents can be performed much faster by this method. Similar to methods described for hemolymph collection (Zanni et al., 2017), centrifugation enables a much faster extraction and investigation of crop contents. However, the two major sources of error - the regurgitation during freezing and contamination from ruptured abdominal tissues releasing other body fluids - may still lead to a significant loss of crop content and reduced analytical accuracy.

Our advanced approach eliminates these major limitations of previously described sampling methods, including loss of crop content, time-consuming manual extraction, limited storage options, contamination with other body fluids and loss of the bee’s body for further analyses.

We here present a novel and simple protocol enabling the experimenter to perform fast and efficient sampling of honey bee crop content through whole bee centrifugation. The composition and the quantity of the crop content can be analyzed reliably without contamination. Samples can be stored for later analysis at two stages of investigation and the body of the bee remains available for further analyses. Furthermore, our protocol may be adapted for other model pollinators such as bumblebees or mason bees.

## Material and Methods

### General procedure

The overall procedure for sampling crop contents is organized into six main steps (**Fig. 1**): **(1)** Sampling of returning foragers, **(2)** Immobilization of sampled bees via cooling, **(3)** Transfer of individual bees into modified tubes, **(4)** Freezing of samples for storage or analysis, **(5)** Centrifugation to extract crop content and **(6)** Analysis of crop contents. A detailed description of each step is provided below. This workflow allows storage of frozen samples at two distinct time points, either after the transfer into the separation tubes or after centrifugation (**Fig. 1**, steps 4 and 5). Consequently, the obtained samples can be analyzed independently at a later stage, enabling the processing of larger sample sizes compared to the immediate analysis required by other methods. To illustrate the advantages of this novel method, we conducted tests on two key pollinator species: the Western honey bee (*Apis mellifera*) and the buff-tailed bumblebee (*Bombus terrestris*). The detailed protocol used in this proof-of-concept is described in the section “Experimental procedure”.

**Figure 1:**
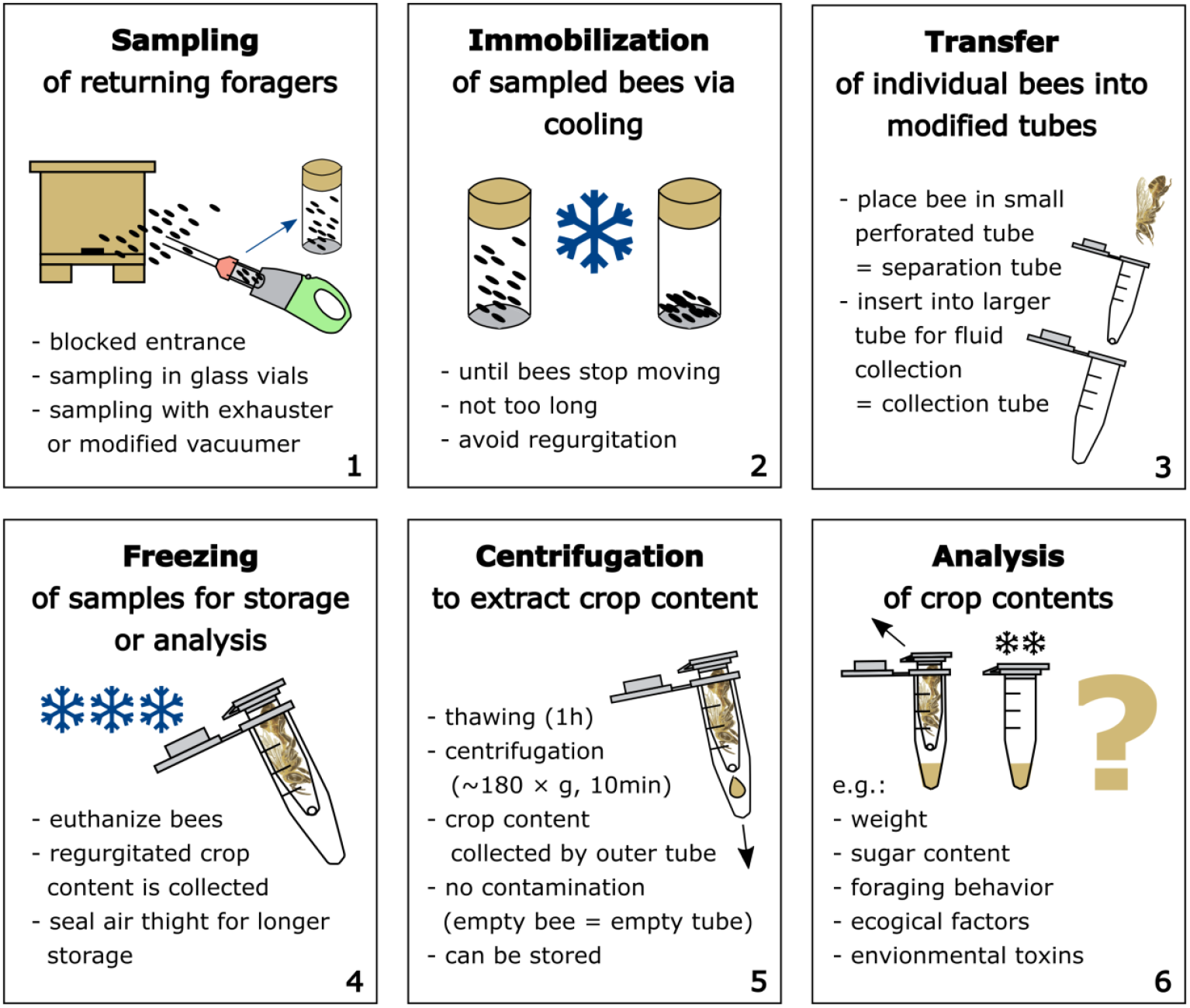
Overview of a rapid and straightforward general procedure for mass sampling of honey bee crop contents in six steps: **(1)** Sampling of returning foragers, **(2)** Immobilization of sampled bees via cooling, **(3)** Transfer of individual bees into modified tubes, **(4)** Freezing of samples for storage or analysis, **(5)** Centrifugation to extract crop content and **(6)** Analysis of crop contents. Samples can be stored in a freezer for later processing either after freezing **(4)** or after centrifugation **(5)**.

#### (1) Sampling of returning foragers

Depending on the study design, it is essential to define which individuals are of interest (e.g., marked bees, returning foragers, nectar foragers, or foragers active or returning within a specific period of time). Individual bees are gently captured in flight by an experienced experimenter or beekeeper using glass vials or aspirators. Mass sampling can be achieved with nets or suction devices, such as a modified handheld vacuum cleaner (**Fig. 1**, Hilsmann et al., 2025). Temporarily blocking the nest entrance can facilitate the capture of returning foragers and reduce the inadvertent collection of newly departing individuals. Scheiner et al. (2013) provides a comprehensive overview of methods for capturing free-flying bees.

#### (2) Immobilization of sampled bees via cooling

After sampling, bees are gently cooled to achieve rapid immobilization. This minimizes the risk of crop content consumption or trophallaxis, particularly during mass sampling, when bees tend to cluster. Ice can be used effectively in the field, while a short exposure to -20 °C in the freezer serves the same purpose in the laboratory. A detailed overview of different immobilization methods is provided by Human et al. (2013). For mass sampling, lower temperatures, such as dry ice or a freezer (-20 °C) are recommended, because clustering individuals generate additional heat. It is essential to remove the bees at the first sign of immobilization, since prolonged exposure can be lethal and may induce regurgitation. Careful monitoring of the cooling process is therefore required, with adjustments to exposure time and temperature based on environmental conditions, container material (e.g., plastic, glass, or metal), and the species or physiological condition of the sampled individuals.

#### (3) Transfer of individual bees into modified tubes

Prior to the experiment, 0.5 ml microcentrifuge tubes (Eppendorf, Germany) are punctured at the lowest point of the tapered end using a metal pin preheated over a Bunsen burner (**Fig. S1**, supplement). Immobilized bees are individually positioned head-first in these so-called separation tubes. Each tube containing a bee is then inserted into a second, 1.5 ml Eppendorf tube, allowing crop contents to be subsequently separated and centrifuged through the small hole of the separation tube into the larger collection tube. For bumblebees, the inner component can be substituted with a cut-off pipette bulb from a disposable 6 ml transfer pipette (Sarstedt, Germany), and 5 ml Eppendorf tubes can be used to collect the fluids (**Fig. S2**, supplement). It is important to prepare these tubes in sufficient numbers beforehand, according to the predetermined sample size. For weight analysis of crop contents, it may be necessary to record the tare weight of the collection tubes if they are not standardized. The principle of the separation tube and the collection tube – an inner smaller vial with hole placed inside a larger outer tube - may be adapted to any insect species by selecting appropriately sized tubes or other centrifugable laboratory plastics.

#### (4) Freezing of samples for storage or analysis

Once placed inside the separation tube and collection tube assembly, samples are immediately transferred to ice or a freezer to euthanize the bees prior to centrifugation. Any crop contents regurgitated during this process are retained entirely for analysis. Long-term storage is possible at this stage, but samples should be sealed airtight to prevent evaporation or freeze-drying, as the lids of the collection tubes cannot be closed when loaded with the separation tubes.

#### (5) Centrifugation to extract crop content

Prior to centrifugation, frozen samples are carefully thawed at room temperature to restore the crop contents to a fully liquid state. The thawing duration should be adjusted according to the size of the insects. During centrifugation, crop contents are expelled through the esophagus and the hole at the tip of the separation tube into the collection tube. As a practical guideline, honey bees can be centrifuged at ∼180 × g for 10 minutes (1,300 rpm; Eppendorf 5424R, Rotor FA-45-30-11, r_max = 9.5 cm) following a thawing period of one hour, whereas bumblebees may require ∼270 × g – 1,700 × g for 10 minutes (1,600 rpm; 4,000 rpm; Eppendorf 5430, Rotor F-35-6-30, r_max = 9.5 cm) after two - three hours of thawing. Because of to their fairly uniform body size, honey bees generally permit a uniform protocol, whereas bumblebees exhibit a size dimorphism that may necessitate adjustments of centrifugation speed according to the individual body size. It is advisable to test all parameters prior to application and adjust them, if necessary, because seasons, climatic/laboratory conditions, or physiological differences within species may have an effect (see “Experimental procedure” for details). Following removal of the separation tube, crop samples can be frozen or analyzed immediately. Furthermore, the emptied bee bodies may be suitable for additional analyses that tolerate a brief interruption of the cold chain. In preliminary tests, we demonstrated a procedure to verify whether the applied centrifugation force is sufficient to expel the entire crop content without contamination. Such contamination is readily identifiable: fresh honey bee hemolymph is initially clear to pale yellow but darkens rapidly upon exposure due to phenoloxidase-driven melanization (Zufelato et al., 2004; Whitten et al., 2017), whereas pollen-rich gut contents typically appear yellow to opaque, reflecting the accumulation of digested pollen material (Brown et al., 2022; **Fig. S3**, supplement).

#### (6) Analysis of crop contents

The weight of the crop contents can be determined by subtracting the tare weight of the collection tube (previously measured or standardized, e.g. for 1.5 ml microcentrifuge tubes) from the weight of the collection tube containing the sample, measured on a laboratory precision balance. Volume measurements can be performed using a laboratory pipette or by determining the height of the liquid in capillaries of known diameter. In our tests, sugar concentration was quantified with a handheld refractometer (RF10-CEI, Extech Instruments, Nashua, USA). Additional analyses of the crop contents may include assessment of impurities, detection of toxins, or identification of pollen residues to verify ingestion of specific substances. These measurements provide a comprehensive characterization of the ingested material and can be adapted according to the aims of the study.

### Experimental procedure

To demonstrate the reliability and applicability of the procedure under experimental conditions, we conducted three main experiments: a proof of concept in honey bees, a method application in bumblebees, and an example of large-scale application in two honey bee subspecies.

As a **proof of concept in honey bees**, we tested whether our method reliably extracted the full crop content and if this was possible without contamination from other body fluids. Bees were initially fed *ad libitum* with a green-colored sucrose solution. After different periods of food deprivation, they were fed a defined volume of unstained sugar solution. The purity of the centrifuged crop contents was assessed by comparing their volume, color and sugar concentration to the ingested solution.

Honey bee foragers were collected at the nest entrance with a modified handheld vacuum cleaner (Hilsmann et al., 2025) **(1)**. The animals were housed in cages and were provided *ad libitum* with a 30% sucrose solution stained with copper chlorophyllin (green, Wusitta, Sitzendorf, Germany). Bees were immobilized by cooling and were mounted in brass tubes for fixation (Scheiner, et al., 2013). After two starvation periods (150 and 180 minutes), the bees were fed 20 µl of pure sucrose solution (30%), were cooled again **(2)**, and were centrifuged using the separation tubes as described above **(3 and 4)**. The centrifugates of both groups were weighed and analyzed with respect to coloration and sugar concentration **(5)**. All centrifugates of sufficient volume were tested for sugar concentration with a handheld refractometer (RF10-CEI, Extech Instruments, Nashua, USA) **(6)**. Crop contents were also inspected visually for signs of contamination from the green solution ingested before starvation and experimental feeding. The presence of green color would indicate an interference in the centrifugation of crop contents by either 1) insufficient starvation (with residual food still in the crop) or 2) contamination due to backflow from the midgut into the crop. If the liquid samples contained hemolymph, they darkened to gray within a few minutes due to oxidation, whereas intestinal contents caused the samples to appear cloudy or yellow (6).

To test our protocol in other pollinators we performed a **method application in bumblebee**s. Buff-tailed bumblebee colonies *(Bombus terrestris*) were purchased from Koppert (Koppert Nederland B. V., Berkel en Rodenrijs, Netherlands). We collected bumblebees directly from study colonies kept in a flight room **(1)**. They were housed in cages and provided *ad libitum* with a 30% sucrose solution stained with copper chlorophyllin (green, Wusitta, Sitzendorf, Germany). Bumblebees were then immobilized by cooling and mounted in 5 ml microcentrifuge tubes (Eppendorf, Hamburg, Germany) modified with a large hole at the tip for subsequent starvation and feeding (**Fig. S4**., supplement). After a starvation period (30 or 120 minutes), the animals were fed pure sucrose solution (30%) until they stopped feeding. The large hole allowed free movement of the bumblebees’ heads for precise individual feeding (see **Fig. S4**, supplement). Since bumblebees did not ingest a predefined volume and varied considerably in intake, the consumed volume was recorded individually. The bumblebees were then cooled again following the protocol **(2)** and inserted into the cut pipette bulb of a 5 ml disposable plastic pipette as described above **(3);** see also **Fig. S2**, supplement) for freezing **(4)** and centrifugation **(5)**. Because bumblebees vary in both body size and food intake (Goulson, 2003), two centrifugation speeds (270 x g and 1,700 x g) were applied. The centrifugates of all groups were analyzed for volume, sugar concentration, and potential contamination as described above **(6)**.

In addition to these initial calibration experiments we used our method in a **large-scale application for two different honey bee subspecies**. We investigated the foraging behavior by analyzing crop content obtained from mass-sampled foragers in the field. Two *Apis mellifera* subspecies were therefore kept in a common apiary (colony setup described in Scheiner et al., 2021). Returning foragers of *A. m. carnica* and

*A. m. ligustica* were collected at their nest entrances as described in **(1)**. After removing and recording any carried pollen loads, the foragers were processed as described above **(2 -5)**. The extracted crop contents were weighed and analyzed for sugar concentration as described in **(6)**. We defined a forager type for each captured individual: water foragers, with crop content <5% sugar concentration; nectar foragers, with crop content >5% sugar concentration and without pollen loads; pollen foragers, carrying a pollen load but no crop content; and mixed foragers, carrying a pollen load together with crop content >5% sugar concentration.

### Statistics

All statistical analyses were performed within the R environment, R version 4.5.1 (R Core Team, 2025). Prior to analysis, the data was tested for normal distribution and homoscedasticity using Shapiro–Wilk and Levene tests, respectively. Since most datasets were not distributed normally, non-parametric tests were applied. For comparisons between two independent groups, Mann-Whitney U tests were applied. We compared crop load weight and sugar concentration between two starvation times (150 min vs. 180 min) in honey bees prior to feeding (**Fig. 2B, C**), as well as between two honey bee subspecies (*A. m. carnica* and *A. m. ligustica*; **Fig. 3B, C**). Differences among more than two treatment groups in combinations of starvation time prior to feeding and centrifugation conditions in bumblebees were analyzed using Kruskal-Wallis tests. Occurrence of contamination in centrifugates was analyzed using Fisher’s exact tests (**Fig. 2A; 4B**). Deviation from initially fed volume and concentration was analyzed using one-sample Wilcoxon tests (**Fig. 2B, C; 4C, D**) in comparison to the respective reference.

**Figure 2:**
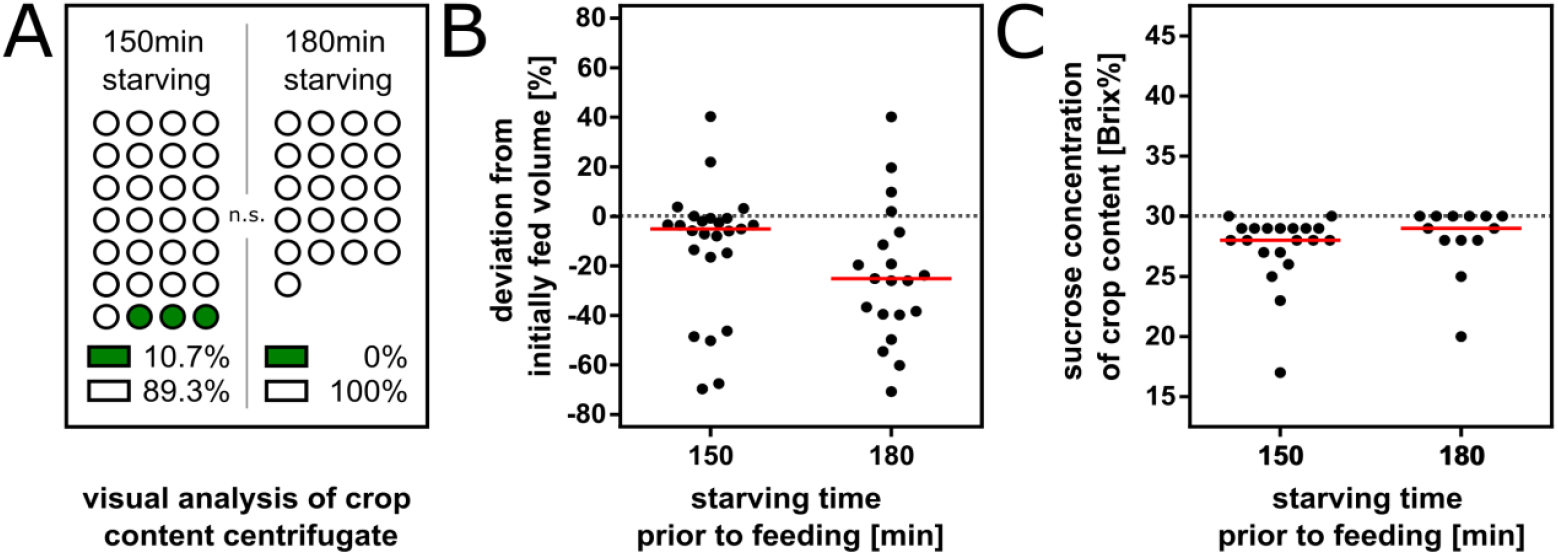
Methodological tests demonstrate that centrifugation of honey bee crop contents results in uncontaminated samples and reliably recovers the initially fed crop contents. Bees were kept in cages and fed *ad libitum* with green-stained sucrose solution. After starvation (150 or 180 min), they were fed 20 µl clear sugar water and processed using the described method. Most bees passed the pre-consumed food (green) to the digestive tract during starvation so that it was not visible in the obtained crop content **(A)**. Green color was only observed in three samples after 150 min starvation but never after 180 min starvation, and the frequency of crop content staining did not differ significantly between starvation times (see **Tab. 1**). The volumes of liquids collected were mostly equal to or smaller than the fed volume **(B)**. While deviations from the initially fed volume were significant within both starvation groups, no significant difference was detected between starvation times (see **Tab. 1**). The concentration of crop contents did not differ between the two deprivation groups and was generally slightly lower than the initially fed solution of 30 % sucrose **(C**; **Tab**.1).

**Figure 3:**
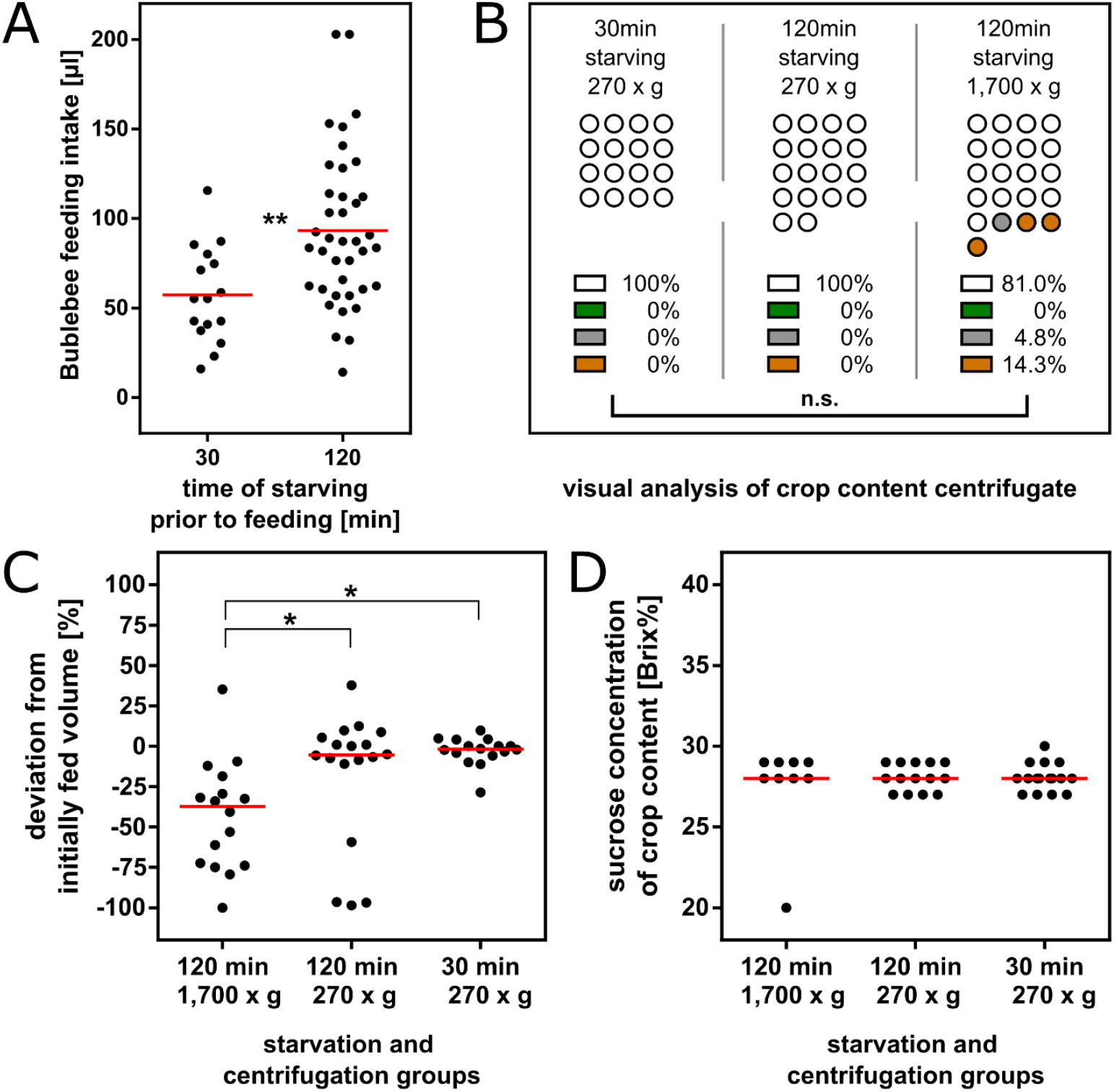
Application of the crop content centrifugation method to buff-tailed bumblebees (*Bombus terrestris*). Because of their larger body size and variable feeding behavior, the protocol required adjustments, but crop contents could still be obtained with minimal contamination. Bumblebees showed a broad range of intake volumes ranging from 14 to 202 µl **(A)**, with individuals starved for 120 min ingesting more sucrose solution than those starved for 30 min (**Tab. 2**). All individuals transferred the previously ingested green solution into the digestive tract during starvation, so no contamination was visible in the crop samples obtained later **(B)**. Only centrifugation at 1,700 × g after 120 min of starvation caused contamination in a few cases. The overall frequency of contamination did not differ significantly among starvation and centrifugation treatments (**Tab. 2**). Because intake volumes varied strongly, the volume of the recovered crop content was expressed as deviation from ingested volume **(C)**. Deviations differed among treatments, with more negative deviations observed after centrifugation at 1,700 × g, and all groups showing deviations below the expected value of zero (**Tab. 2**). Sugar concentrations of crop contents were similar across treatments and slightly lower than the fed concentration of 30 Brix% **(D;** see **Table 2**).

## Results

### Proof of concept in honey bees

Bees were initially fed *ad libitum* with a green-colored sucrose solution. Subsequently, they were starved for 150 or 180 minutes before being fed with a defined volume of clear 20 µl sucrose solution. Among the subsequently collected crop contents, all 21 samples collected after 180 minutes starvation appeared clear without visible contamination. In contrast, crop contents from bees deprived for 150 minutes contained three green-colored samples out of 28 (12.5%). This difference in contamination was not statistically significant (**Fig. 2A**; **Tab. 1**).

**Table 1.**
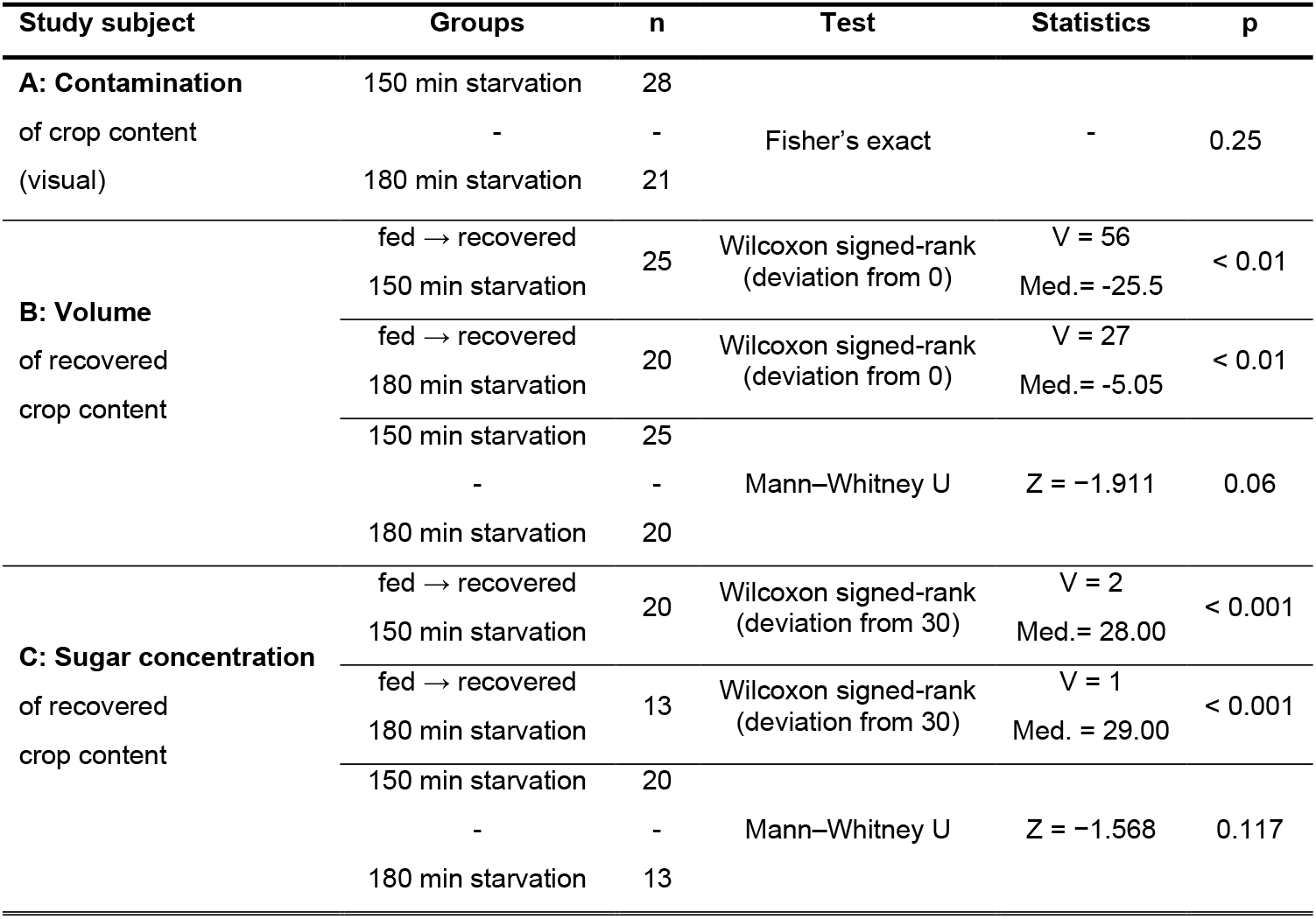
Test statistics for comparisons of **(A)** Statistical summary of tests performed to validate the centrifugation-based method for isolating crop contents in honey bees after two different starvation times (150 min and 180 min). Analyses compare crop content contamination frequency after starvation, **(B)** deviations between the initially fed and recovered crop volume, and **(C)** sugar concentration of recovered crop contents relative to the initially fed sucrose solution in honey bees starved for 150 minutes or 180 minutes prior to investigation. See Fig. 2 for graphical display.

**Table 2.**
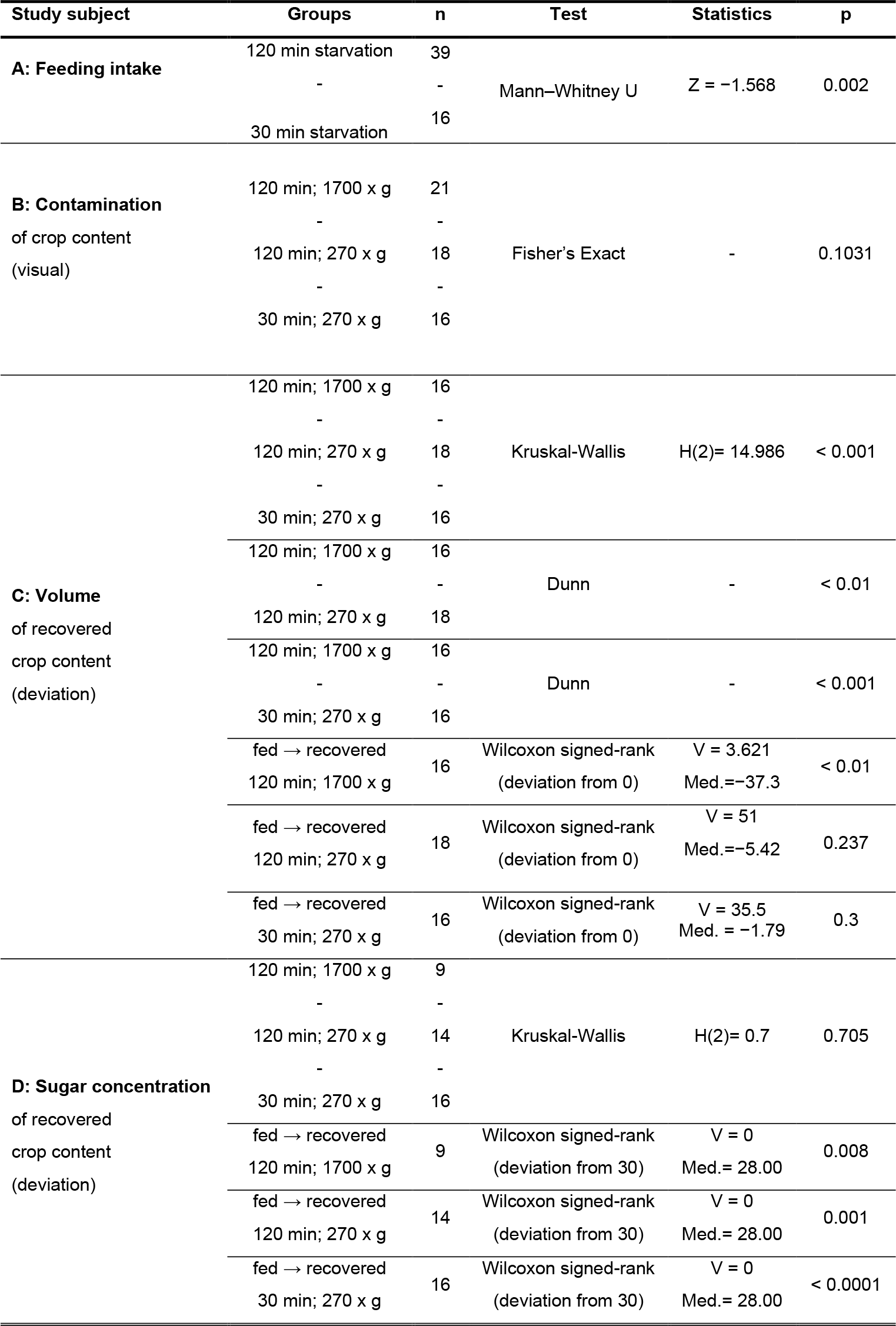
Test statistics for comparisons of **(A)** intake volumes after different starvation times (30 min and 120 min), **(B)** crop content contamination frequency across starvation and centrifugation treatments, **(C)** deviations between the initially ingested and recovered crop volume, and **(D)** sugar concentration of recovered crop contents relative to the initially fed sucrose solution in buff-tailed bumblebees (*Bombus terrestris*). See **Fig. 3** for graphical display.

Values of crop volume deviations tended to be smaller in bees food-deprived for 180 minutes compared to those food-deprived for 150 minutes *(p = 0*.*06*, **Fig. 2B**; **Tab. 1)**. However, the volumes in both groups were generally slightly smaller than the 20 µl initially fed and thus showed significant deviations from the initial volume (0% deviation), respectively (**Fig. 2B**; **Tab. 1**).

Sugar concentrations did not differ between the two deprivation groups but were generally lower than the initially fed solution of 30% sucrose (**Fig. 2C, Tab. 1**).

### Method application in bumblebees

To assess the suitability of our method for bumblebees, we analyzed crop contents from buff-tailed bumblebees (*Bombus terrestris*) that were food-deprived for 30 or 120 minutes after an *ad libitum* diet of green-stained sucrose solution. After food deprivation, bumblebees were fed with clear sucrose solution until they stopped intake.

Bumblebees showed a highly variable intake volume (14-202 µl), with significantly higher intake volumes after a longer starvation period of 120 min compared to 30 min (**Fig. 3A, Tab. 2**). No green contamination was detected in the crop samples, indicating that all individuals from the three starvation groups cleared the green-stained solution consumed before starvation into the digestive tract (**Fig. 3B**). Only centrifugation at 1,700 x g after 120 min of starvation caused contamination with body fluid in a few cases: one sample with hemolymph (4.8%) and three with yellow opaque gut contents (14.3%). The overall frequency of contamination did not differ significantly across differently treated bumble bees (starvation time and centrifugation force, **Tab. 2**).

Intake volumes also varied strongly among individuals of the same starvation group. Thus, extracted crop content volume was expressed as a deviation from the initially ingested volume on individual basis (**Fig. 3C**). The four obviously contaminated samples (hemolymph and gut contents) showed between 47% and 157% more volume than were initially fed and were excluded from further analyses. Deviations differed significantly among treatments. Significantly larger negative deviations (i.e., less volume centrifuged than initially fed) were detected in the group centrifuged with 1,700 x g in comparison to both other groups. Based on these deviations, the centrifuged volume also differed significantly from the initially fed volume (0% deviation) in the group centrifuged with 1,700 x g (**Fig. 3C; Tab. 2**).

Sugar concentrations of crop contents were similar across all starvation and centrifugation treatments and differed significantly from the initially fed concentration of 30% Brix (**Fig. 3D**; **Tab. 2**).

### Large-scale application for two different honey bee subspecies

To test the method under field-realistic conditions and with large sample sizes, we compared the foraging behavior of two honey bee subspecies by analyzing mass-sampled honey bee foragers. Forager types as well as the weight and sugar concentration of their crop contents were examined in 269 individuals of *Apis mellifera carnica* and 322 individuals of *Apis mellifera ligustica*. Pollen-carrying bees were identified, and the crop contents of all captured bees were isolated using the method described above.

Among *Apis mellifera carnica* foragers, 83 individuals (30.9%) had empty crops, 50 (18.6%) carried pollen only, 74 (27.5%) carried nectar only (sugar concentration >5%), 49 (18.2%) carried both pollen and nectar, and 13 (4.8%) carried water (sugar concentration <5%). Corresponding values for *A. m. ligustica* were 122 (37.9%) empty foragers, 85 (26.4%) pollen foragers, 72 (22.4%) nectar foragers, 30 (9.3%) foragers of pollen and nectar, and 13 (4.0%) water foragers. These proportions differed significantly between subspecies (**Fig. 4A, Tab. 3**).

**Table 3.**
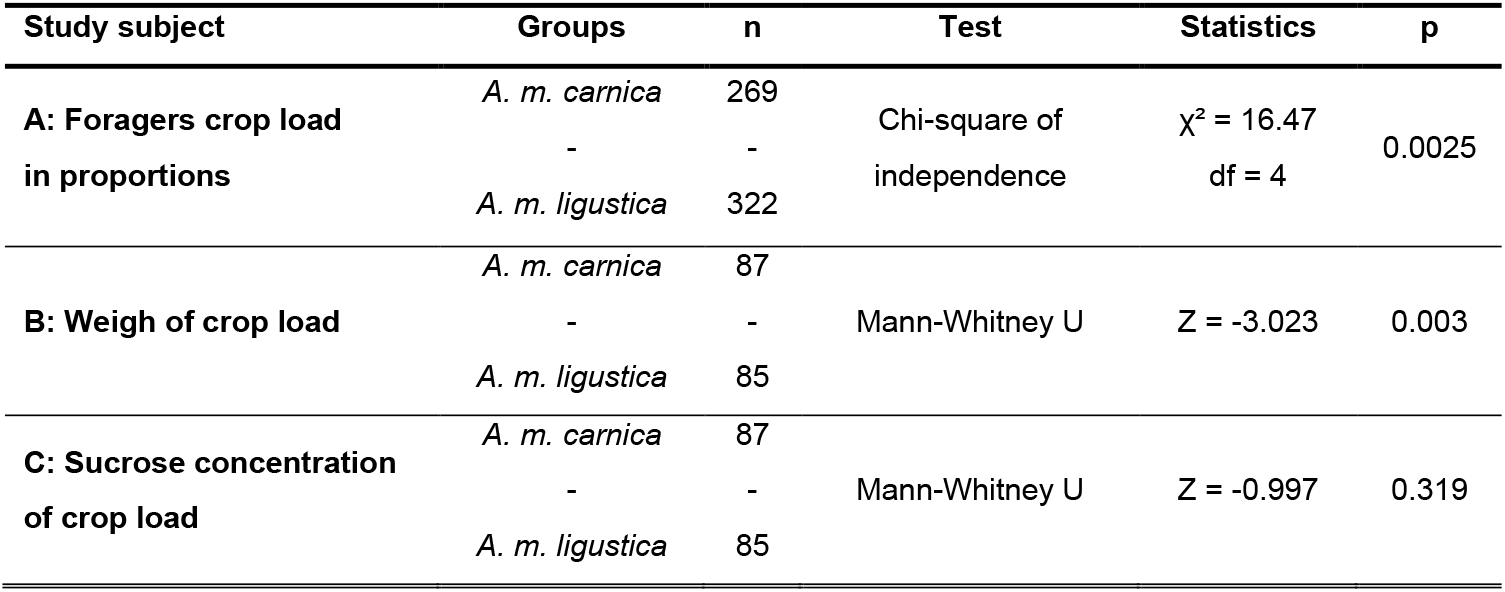
Test statistics for comparisons of **(A)** the distribution of forager types between subspecies, **(B)** crop load weight of pollen-free foragers, and **(C)** sugar concentration of crop contents in pollen-free foragers in mass-sampled honey bee foragers from two subspecies (*Apis mellifera carnica* and *Apis mellifera ligustica*). See **Fig. 4** for graphical display.

**Figure 4:**
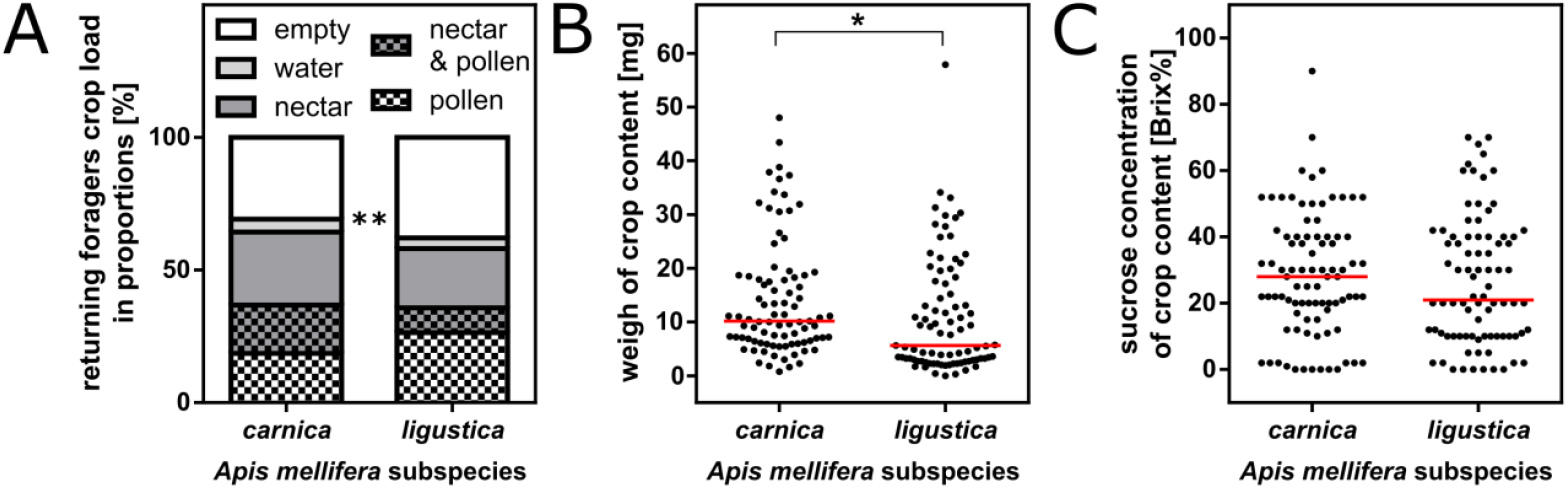
Large-scale application of the centrifugation method to compare crop contents of mass-sampled honey bee foragers from two subspecies (*Apis mellifera carnica* and *Apis mellifera ligustica*) under field-realistic conditions. Foragers were classified by type (empty, water forager, nectar forager, mixed nectar–pollen forager, and pollen forager), and the weight and sugar concentration of their crop contents were measured. Among 269 *A. m. carnica* and 322 *A. m. ligustica foragers*, the proportions of forager types differed significantly between subspecies **(A)**. To compare crop loads independent of pollen transport, only pollen-free foragers were considered. Crop load weights differed between subspecies **(B)**, whereas sugar concentrations of the crop contents did not **(C)**. Statistical analyses are summarized in **Tab. 3**.

Comparing the crop load weights of 87 *A. m. carnica* and 85 *A. m. ligustica* foragers without pollen, *A. m. carnica* foragers carried significantly higher crop loads **(Fig. 4B, Tab. 3**). In contrast, the sugar concentrations of the crop contents collected by these foragers did not differ significantly between subspecies **(Fig. 4C, Tab. 3**)

## Discussion

### Proof of concept in honey bees

The newly described centrifugation method effectively isolates the pure crop load of honey bees without contamination from other parts of the digestive system. Previously ingested colored food was never encountered in the centrifugate after 180 minutes of starvation and occurred only occasionally after 150 minutes of starvation. Likewise, the volume of the centrifugate was equal to or slightly smaller than the initially fed volume. Both indicate that no fluids are expelled during centrifuging once they have passed the honey crop. This reflects the honey bee anatomy: a muscularly controlled proventriculus acts as a valve between the crop and the midgut, regulating the passage of fluids and preventing backflow (Blatt & Roces, 2002; French et al., 2014).

We obtained smaller crop content volumes from bees that had been food-deprived for a longer period, which additionally indicates the safe separation of crop content from other body fluids. Since both groups were cooled immediately after feeding, these animals must have transferred the sucrose solution faster to the midgut than the less food-deprived group. The reduced volume in the longer starved group therefore suggests that even fluids, transported from the crop to the midgut shortly before freezing, are not extracted by our method. This observation also aligns with reports of variable retention times of nectar in the honey crop, depending on e.g., the bee’s activity and energy demand (Nicolson & Thornburg, 2022, Crailsheim, 1988). Accordingly, the rare occasions of green traces in the centrifugates of shortly food-deprived bees indicate that some of those bees retained the stained solution in their crop for a longer period. For future experiments, it is therefore essential to plan precisely when, where and how the animals are collected and processed to ensure that the crop content represents the physiological state at that specific time.

Additionally, it is crucial to freeze the insects as soon as possible after collection and transfer to the separation tube to prevent further consumption or processing of crop content.

We did not detect any visible contamination due to tissue ruptures before determining the sugar concentration of the crop content. Such damages would have been easily notable in the clear centrifugate since hemolymph oxidizes and turns gray while pollen-rich gut contents appear yellow or opaque in solution (Zufelato et al., 2004; Whitten et al., 2017; Brown et al., 2022). Thus, the centrifugation method and force (∼180 × g for 10 minutes) tested seems suitable for crop content extraction without damage to intestinal tissues. Although a dilution by body fluids seems unlikely, the sugar concentration of crop contents was slightly lower than in the initially fed solution. Even though sugar absorption mainly happens in the midgut (Crailsheim, 1988), earlier metabolic processes thus seem to have reduced the sugar concentrations of the crop content. Such metabolic activity within the crop has been associated with the presence of enzymes and signaling molecules influencing sugar conversion and fluid processing (Kodrík et al., 2022; Pavlović et al., 2025). Similarly, like the volume of crop contents, their sugar concentration may be influenced by various factors that affect food digestion. For example, temperature, feeding history and age can affect digestion rate and nutrient transfer between crop and midgut considerably in honey bees (Crailsheim, 1990; Crailsheim, 1992). We cannot further specify which dilutions or digestive processes may have caused the reduced concentrations observed. However, under field-realistic sampling conditions these factors reflect the real-world situation given the standardized and uniform handling of all individuals. When collecting samples, these factors should thus be considered and harmonized to ensure comparable results. In addition, centrifugates should be checked visually for spilled hemolymph (grey) or gut contents (yellow/opaque) based on their coloration (Zufelato et al., 2004; Whitten et al., 2017; Brown et al., 2022) before further investigation, which can easily be implemented in the sample processing.

### Method application in bumblebees

Examining crop content in bumblebees is more complex than in honey bees. Bumblebees display a higher diversity, both in their body size and in their behavioral roles within the colony (Holland et al., 2021; Fitzgerald et al., 2022). This diversity is also reflected in the strongly differing intake volumes observed in our feeding experiment. Nevertheless, the intake levels were in general higher after longer periods of food deprivation, which fits well to the general pattern discussed for honey bees above. This should be kept in mind when handling animals in the field or studying individual bumble bees in laboratory setups. Because of the wide variations in intake volumes, we compared the percentage deviation between the recovered crop contents (centrifugate) and the amount of sugar solution fed initially.

Regardless of the size differences, the centrifugation force of 270 x g proved well for the crop content extraction of bumblebees. Crop contents could be retrieved without significant deviations from the initially fed volume using this method. The centrifugation force thus appeared strong enough to extract crop contents through the esophagus of the tested bumblebees. On the other hand, the procedure did not result in contamination from other body fluids or stained sucrose solution ingested before experimental feeding. The centrifugation force of 270 x g apparently did not damage the intestinal tissue or the proventriculus (i.e., valve between crop and midgut that regulates fluid flow; Hüftlein et al., 2024) of test animals. In contrast, the stronger centrifugation force of 1,700 x g resulted in both contamination from other body fluids and significantly lower recovered volumes. Both effects might be caused by tissue rupture and liquid loss within the abdominal lumen. Likewise, a higher centrifugation force may compress the esophagus in a way that reduces its permeability for crop contents. Depending on the type and size of examined bumblebees (e.g., smaller foragers or different species), centrifugation speed should thus be adjusted based on these findings and validated through appropriate pre-tests for each species and study setup.

In addition to the generally higher initial intake, a longer starvation time also led to greater variation in the crop content volumes retrieved after centrifugation. As with honey bees, this could indicate either a faster absorption of the fed sucrose solution into the midgut or an early-stage digestion by, for example, glandular secretions within the crop (Carnell et al., 2020). This again demonstrates that our method represents a snapshot of the field-realistic crop content status at the time of freezing and emphasizes the importance of appropriate methods for rapid field sampling. In any case, samples should be frozen as quickly as possible after collection and separation of bees so that the animals do not starve, metabolize the crop content or suffer from the stress of prolonged handling.

Similar to the slightly decreased sucrose concentration found in honey bee crop centrifugates sucrose concentration of bumble bee crop contents was overall decreased by 2% from the initially fed 30% sucrose solution. Thereby, the concentration did not differ between the groups in terms of starvation period (30 min and 120 min), centrifugation force (270 x g and 1,700 x g) or both factors combined. As in honey bees, this points towards a general metabolic decrease of sucrose concentration (i.e., field-realistic condition) rather than an effect of the method itself. We consider the slight dilution of the liquid to be part of the natural metabolic processes occurring within the crop, which are mediated by internal enzymes such as α-glucosidases within the crop and midgut walls (Pavlović et al., 2025), and thus an inherent part of the foraging process as described above for honey bees. The experiments with bumblebees demonstrate that the method can be adapted and applied for non-Apis pollinators. Indeed, many nectar-feeding hymenopteran taxa — including commonly studied bees of the genera *Megachile, Osmia, Xylocopa* and *Halictus* — possess a crop and proventriculus anatomy which appears sufficiently similar to allow an application of this method (Ruiz-González et al., 2022). However, the methodology needs to be adapted to the respective species and research questions by means of suitable extraction materials and centrifugation parameters. The potential realm of application might thus extend to dozens (if not hundreds) of nectar-collecting insect species and thus enable insights into finely ramified niches of foraging ecology.

### Large-scale application for two different honey bee subspecies

Our mass sampling experiment in the field provides a clear example of a practical application of the method in pollinator research. In addition, it adds value to the initial proof of concept. The occurrence of “empty bees” (i.e., no pollen collected and no fluid retrieved after centrifugation in the collection tube) in both subspecies confirms the above-mentioned findings that neither pre-consumed food from the midgut nor other body fluids are retrieved in the centrifugate. Vice versa, these free flying bees appeared to be truly “empty”, since the initial method of evaluation proved that crop contents of experimentally fed animals could be retrieved in all cases (i.e., fed bees with full honey crops were never incorrectly classified as “empty”).

Since sampling took place at forenoon, before orientation flights occur (Capaldi & Dyer, 1999), the high portions of empty bees indicated a temporary shortage of nectar-flow at the time of sampling. This phenomenon is consistent with the strong temporal and environmental variability of floral nectar reward (Plos et al., 2023; Parachnowitsch & Kessler, 2018). The proportion of water foragers was similar in both subspecies, whereas the proportions of nectar foragers, pollen foragers and mixed nectar-pollen foragers differed. These differences in the composition of foraging force seem to be directly related to subspecies affiliation as described for other traits before (Uzunov, et al., 2014; Scheiner, et al., 2021) and as was hypothesized for foraging behavior in particular (Cakmak et al. 2010).

Examining the crop contents more closely, the *A*.*m. ligustica* foragers transported significantly smaller crop loads compared to *A*.*m. carnica*, while both subspecies foraged on nectar resources of similar concentration. The weight of crop contents investigated was low overall compared to those reported in former studies (Seeley, 2014; De la Barrera & Nobe, 2004; Hunt, et al., 1995; Eckert, et al., 1994, Park 1926). This again could reflect poor overall foraging conditions, i.e., scarce nectar flows during summer (Hunt, et al., 1995; Waser & Price, 2016; Wyatt, et al., 1992). Nevertheless, the relative differences between crop loads of both subspecies investigated points towards a better adaptation to those foraging conditions in the locally derived *A*.*m. carnica* stock compared to the *A*.*m. ligustica* colonies introduced for this study. Indeed, previous physiological studies suggest that *A. m. carnica* exhibits greater efficiency under central European temperate conditions, whereas *A. m. ligustica* may perform better under other climatic regimes (Kovac et al., 2014).

## Conclusion

Overall, our novel method provides a straightforward, reliable and efficient protocol for studies in individual foraging behavior as well as for rapid large-scale sampling of honey bee crop contents. It thus opens up new realms of application for honey bees as bioindicators and model organisms.

The described method yields comparable results, circumvents the problem of regurgitation and substantially reduces processing time. Our proof of concept furthermore highlights important considerations for the calibration of the experimental parameters before applying the technique in individual research questions.

In addition, sample processing can be safely interrupted at two stages - either before centrifugation (by freezing the separated individuals) or after centrifugation (by freezing the received crop content as centrifugate), greatly enhancing flexibility for large sample sizes and field studies. Based on our results in bumblebees, we further demonstrate the method’s applicability to other pollinator taxa, emphasizing its potential as a universal tool for comparative studies on nectar foraging, environmental exposure, and pollinator health.

## Acknowledgments

We thank Markus Thamm for his expertise in molecular laboratory work, and Kayun Lim and Stefanie Schmidt for their assistance. We also thank Dirk Ahrens for beehive maintenance.

## Author Contributions

L.D. and M.G. developed the idea underlying the approach. L.D., M.G., and R.S. designed the study, drafted the manuscript and revised the manuscript. J.S. conducted all experiments. J.S. and M.G. performed the data analysis. L.D. and M.G. contributed equally to the idea and this work.

## Availability of data and materials

Correspondence and requests for materials not included in the supplementary information should be addressed to M.G..

## Ethics approval and consent to participate

No ethics approval or consent to participate was required for this study.

## Competing interests

The authors declare that they have no competing interests.

## Notes

### Competing Interest Statement

The authors have declared no competing interest.

