## Supplementary Information: Figures S1,S2,S3 and S4 for "A fast and easy mass sampling method for social bee crop content"

For:

**The following supporting information is available for the article:**

**Figure S1: Equipment needed for crop content centrifugation: a) separation tubes are prepared by piercing a red-hot metal pin through the tapered end of a 0.5 ml Eppendorf tube. b) orientation of the bee within the separation tube (inserted head first). c) separation tube with bee inside the collection tube (1.5 ml Eppendorf tube).**

**Figure S2: Separation tubes for bumblebees were prepared by cutting out the lower part of the bulb from a disposable 6 ml Sarstedt pipette (left). This funnel-shaped plastic piece was then inserted in a 5 ml Eppendorf tube to hold back the bumblebee body (right).**

**Figure S3: Centrifuges contaminated with green stain from trial feeding (left) and hemolymph and green stain (middle), as well as without contamination (right).**

**Figure S4: Feeding of an isolated bumblebee restrained in a 5 ml Eppendorf tube. The tapered end of the tube was cut open to allow feeding while still holding back the bumblebee**

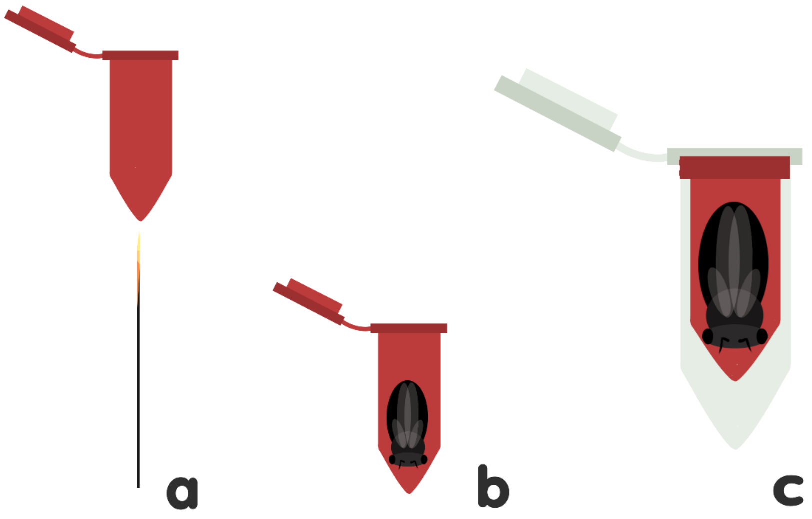


**Figure S1: Equipment needed for crop content centrifugation:** a) separation tubes are prepared by piercing a red-hot metal pin through the tapered end of a 0.5 ml Eppendorf tube. b) orientation of the bee within the separation tube (inserted head first). c) separation tube with bee inside the collection tube (1.5 ml Eppendorf tube).

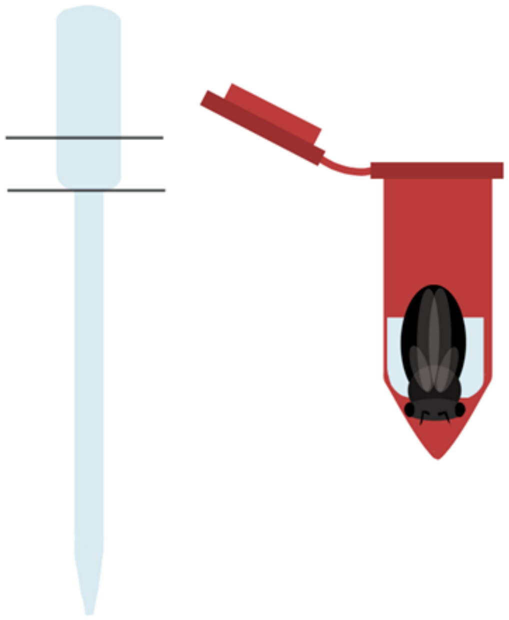

**Figure S2: Separation tubes for bumblebees were prepared by cutting out the lower part of the bulb from a disposable 6 ml Sarstedt pipette** (left). This funnel-shaped plastic piece was then inserted in a 5 ml Eppendorf tube to hold back the bumblebee body (right).

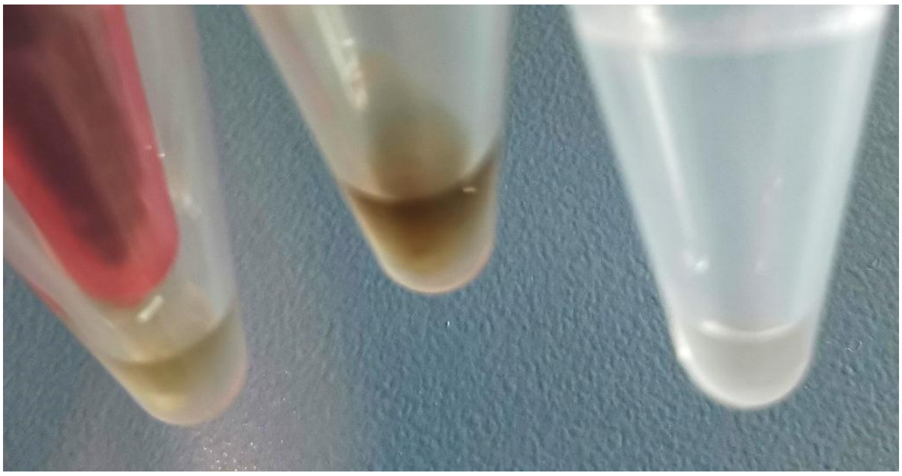

**Figure S3: Centrifuges contaminated with green stain from trial feeding (left) and hemolymph and green stain (middle), as well as without contamination (right).**

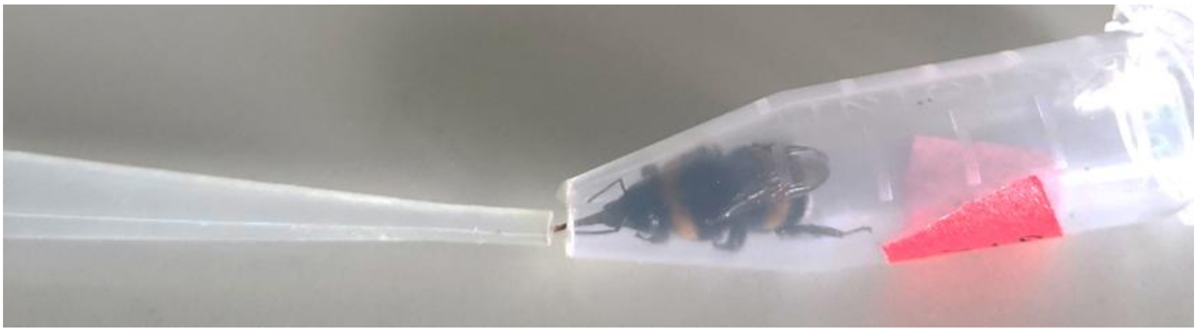

**Figure S4: Feeding of an isolated bumblebee restrained in a 5 ml Eppendorf tube.** The tapered end of the tube was cut open to allow feeding while still holding back the bumblebee
